# Exploring the ability of the MD+FoldX method to predict SARS-CoV-2 antibody escape mutations using large-scale data

**DOI:** 10.1101/2024.05.22.595230

**Authors:** L. América Chi, Jonathan E. Barnes, Jagdish Suresh Patel, F. Marty Ytreberg

## Abstract

Antibody escape mutations pose a significant challenge to the effectiveness of vaccines and antibody-based therapies. The ability to predict these escape mutations with computer simulations would allow us to detect threats early and develop effective countermeasures, but a lack of large-scale experimental data has hampered the validation of these calculations. In this study, we evaluate the ability of the MD+FoldX molecular modeling method to predict escape mutations by leveraging a large deep mutational scanning dataset, focusing on the SARS-CoV-2 receptor binding domain. Our results show a positive correlation between predicted and experimental data, indicating that mutations with reduced predicted binding affinity correlate moderately with higher experimental escape fractions. We also demonstrate that better performance can be achieved using affinity cutoffs tailored to distinct antibody-antigen interactions rather than a one-size-fits-all approach. We find that 70% of the systems surpass the 50% precision mark, and demonstrate success in identifying mutations present in significant variants of concern and variants of interest. Despite promising results for some systems, our study highlights the challenges in comparing predicted and experimental values. It also emphasizes the need for new binding affinity methods with improved accuracy that are fast enough to estimate hundreds to thousands of antibody-antigen binding affinities.

## Introduction

The global impact caused by the Severe Acute Respiratory Syndrome Coronavirus 2 (SARS-CoV-2), the virus responsible for COVID-19, serves as a stark reminder of the need for robust preparedness and research to effectively combat present and future infectious threats. One crucial area of focus is the study of neutralizing antibodies (nAbs) to combat viral diseases^1–4^. These nAbs render viruses noninfectious by binding to functional molecules, blocking viral entry, and ultimately preventing host cell invasion^5^. By May 2022, the pivotal role of nAbs took center stage as 11 monoclonal nAb treatments received emergency use authorization for COVID-19, marking a crucial advancement in managing the pandemic^6^. Additionally, the exploration of numerous nAbs in preclinical or clinical trials underscores their potential as both preventive and therapeutic options against SARS-CoV-2^7^.

Despite the success of Ab-based therapies, the rapid evolution of viruses allows them to develop resistance mutations to nAbs, effectively evading the immune system’s ability to recognize and neutralize the threat (Figure 1a)^8–10^. Experimental work has been done to map mutations in the SARS-CoV-2 spike (S) protein that show antibody escape, with a focus on the spike receptor binding domain (S-RBD). The S-RBD is of particular interest as it serves as the primary binding region for Abs, with S-RBD targeting Abs categorized into four classes based on their targeted structural epitopes (Figure 1b)^1,11,12^. Notably, Starr et al. conducted an experimental mapping of all single amino acid mutations within the S-RBD to determine the impact of each on biophysical properties like Ab binding^13–15^. Their large-scale studies identified mutations and sites associated with Ab-escape using a yeast-displayed deep mutational scanning (DMS) technique. The identification and understanding of Ab-escape mutations, particularly within the S-RBD, is essential for developing effective strategies against viral resistance and for preparing for future pandemics.

**Figure 1.**
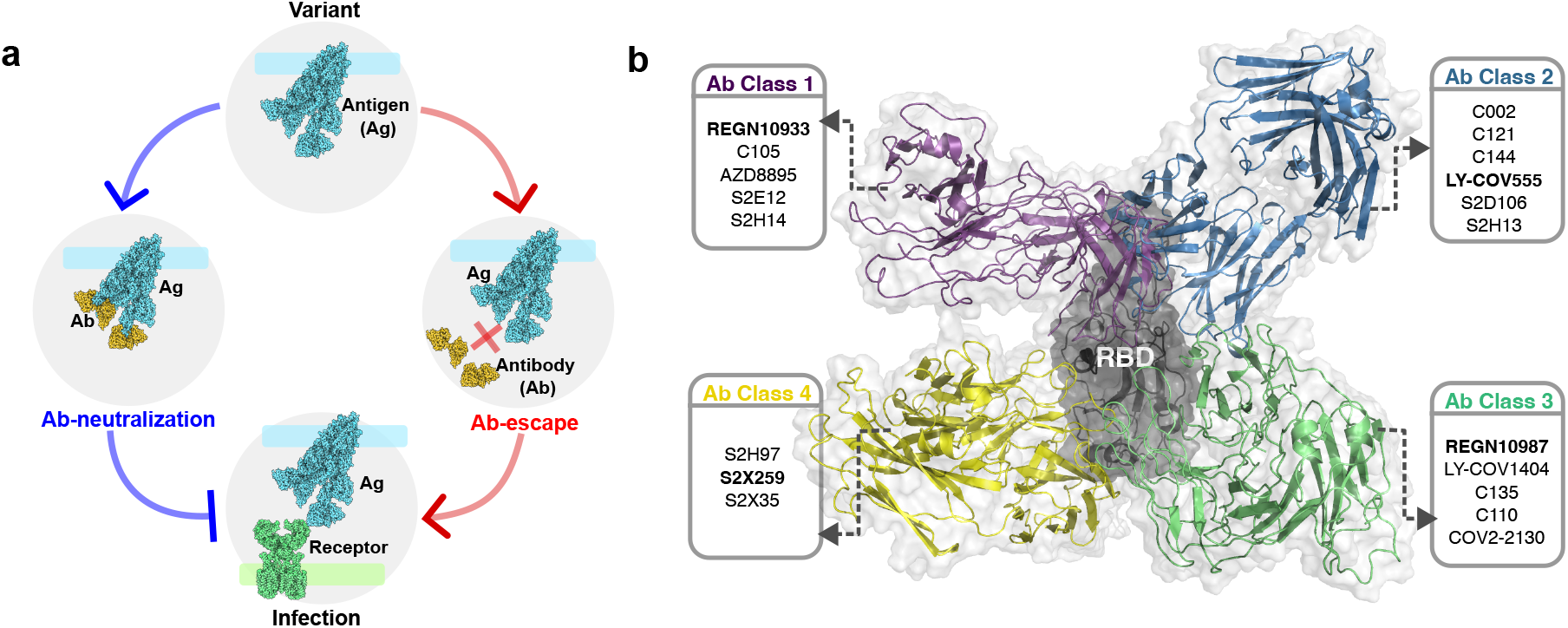
SARS-CoV-2 mutations could lead to Ab escape. a) A wild-type viral antigen (Ag) can mutate into a variant that may either retain Ab-neutralization hence preventing infection, or lead to Ab-escape hence maintaining infection. b) RBD-directed neutralizing Abs can be divided into four main classes based on the epitopes they target^11^. Abs tested in this study are listed according to their class. For each class, one representative Ab bound to the RBD is shown: Class 1, REGN10933; Class 2, LY-CoV555; Class 3, REGN10987; Class 4, S2X259. Protein images on the left were generated with Protein Imager software^16^.

Computational methods for predicting Ab-escape mutation can significantly reduce the time and financial cost compared to experiments, however, they come with notable challenges. First, validation is difficult due to the limited availability of large datasets. Second, the parameters associated with these methods may lack universal applicability. Third, while some methods provide fast results, their accuracy may be compromised, and more accurate methods may be computationally intensive and time-consuming. Current methods range from sequence-based^17,18^ to structure-based^19–22^, including hybrid models^23^. Structure-based approaches are particularly important since protein structure profoundly influences their biophysical properties.

In these methods, force fields are employed to assess the change in the protein binding affinity due to the mutation, aiming to identify destabilizing mutations (i.e., mutations that disrupt binding) that could lead to Ab-escape^24–26^. These force fields can be developed using rigorous yet computationally expensive physics-based potentials, or using computationally less expensive but comparatively less rigorous high-throughput semi-empirical/statistical potentials^27^. Computational work has been done to map mutations in the SARS-CoV-2 S-RBD that show antibody escape by using a structure-based non-rigorous approach. For example, Keng-Chang et al. conducted a computational DMS using multiple semi-empirical/statistical potential-based methods to identify immune-escaping hotspots in the SARS-CoV-2 RBD in complex with five different Abs^20^. S. Huang et al. proposed a hybrid structure/sequence-based model to predict similar high-risk immune-escaping hotspots on the RBD averaging results from 145 non-specified Abs^19^. Barnes et al. used physics-based molecular dynamics (MD) in combination with FoldX to develop a watchlist for pandemic surveillance for the SARS-CoV-2 RBD in complex with Abs B38 and CB6^21^. These studies offer valuable insights, however, there are some reasons to expand on this body of work. First, some of these studies predominantly concentrate on site-level immune-escaping hotspots^19,20^. Second, some do not employ conformational sampling of the experimental structures for their predictions^19,20^. Third, some are limited in their focus on a few specific systems^21,22^. Finally, some present average effects rather than antibody-specific data; this is important since Abs targeting structurally similar epitopes may elicit different responses to the same mutation^19^.

Among the array of possible computational approaches, the MD+FoldX method stands out as a promising avenue for predicting Ab-escape mutations, as evidenced by our previous comparison of eight methods across various protein-protein systems^27^. The MD+FoldX approach was developed by our lab in 2016 and has since been applied to various studies^21,27–30^. It is a 3-D structure-based approach that integrates MD simulations with FoldX estimations. For this method, a set of snapshots of the proteins are generated from atomistic MD simulations, FoldX^31^ is used to estimate binding affinities for each snapshot which are then averaged to generate a single binding affinity estimate per variant, where each variant differs in a single point mutation. These binding affinities can then be used to identify mutations that disrupt binding, typically indicated by high positive values, hence determining possible Ab-escape mutations. This approach holds significant potential as it offers detailed immune-escaping insights at the mutation level, explores diverse protein conformations beyond single experimental structures, and demonstrates versatility in its applicability to a wide range of Abs.

In this study, we capitalize on the wealth of data generated during the COVID-19 pandemic by focusing on the SARS-CoV-2 RBD to tackle significant challenges associated with the MD+FoldX approach. We investigate the validity of using MD+FoldX to predict Ab-escape by comparing to experimental DMS and we examine the generalizability of the energy cutoff that is crucial for identifying escape mutations. Initially, we establish the correlation between MD+FoldX predictions and DMS escape fraction data. Subsequently, we refine the binding affinity cutoff to classify escape mutations and assess the method’s accuracy in identifying escape mutations. Finally, we present the potential escape mutation list generated by our method and propose avenues for future research.

## Methods

In this study, we assessed the predictive capability of the MD+FoldX method for identifying SARS-CoV-2 Ab-escape mutations and examined their transferability across diverse systems using DMS-derived escape fraction data. Our strategy involved selecting SARS-CoV-2 RBD-targeting Abs from four different classes, guided by the availability of experimentally determined complex structures and DMS-derived escape fraction data (Figure 1a). Binding affinity predictions were conducted through the MD + FoldX approach (Figure 1b). Binding affinity predictions were correlated with experimental escape fraction data from DMS experiments (Figure 1c). Subsequently, we refined the affinity cutoff, crucial for selecting predicted escape mutations, tailored individually for each system. The use of such system-specific optimized cutoffs provided precision-based performance and Ab-escape lists.

### 0.1 Data curation

We selected 19 SARS-CoV-2 RBD-targeting antibodies that have experimentally determined Ab/Ag structures and escape fractions from DMS experiments available (Figure 1b and Table 1). Experimental structures had resolution ranges of 1.83-3.0 Å from X-ray crystallography and 3.4-3.95 Å from cryo-EM. We named each Ab/Ag pair after the Ab, given the antigen’s consistency across all pairs. For example, the S2H14 Ab/S-RBD complex is referred to as the S2H14 system. The experimentally determined structures of the Ab/Ag complexes were retrieved from the Protein Databank (PDB) and their corresponding PDB identifiers (IDs) are listed in Table 1. DMS data comes from the yeast-display DMS approach, developed by the Bloom lab^13^ (Refer to Table 1 for DMS data citations). Briefly, it involves creating libraries of all possible monomeric S-RBD single amino acid mutants (Wuhan-Hu-1, Genbank accession number MN908947, residues N331-T531) and filtering out the mutants with significantly reduced ACE2 affinity or folding efficiency. The resulting library is expressed on yeast surfaces and the effect of each mutation on Ab binding is quantified by fluorescence-activated cell sorting and deep sequencing. The escape fraction resulting from each RBD mutation signifies the ratio of cells with that mutation found in the corresponding Ab-escape cell sorting bin. These fractions can range from 0 (indicating no cells with this mutation were found in the Ab-escape bin) to 1.0 (indicating that all cells with that mutation were found in the Ab-escape bin).

**Table 1.** List of antibodies considered in the present study. Each Ab is presented along with its DMS data citation and Ab/Ag co-crystallized structure PDB ID. Ab class is according to the classification scheme of Barnes et al.^11^.

| Class | Ab | PDB ID |
| --- | --- | --- |
| 1 | REGN10933<sup>32</sup> | 6XDG |
|  | C105<sup>33</sup> | 6XCM |
|  | AZD8895<sup>14</sup> | 7L7D |
|  | S2E12<sup>34</sup> | 7K45 |
|  | S2H14<sup>34</sup> | 7JX3 |
| 2 | C002<sup>33</sup> | 7K8T |
|  | C121<sup>33</sup> | 7K8X |
|  | C144<sup>33</sup> | 7K90 |
|  | LY-CoV555<sup>35</sup> | 7KMG |
|  | S2D106<sup>34</sup> | 7R7N |
|  | S2H13<sup>34</sup> | 7JV2 |
| 3 | REGN10987<sup>32</sup> | 6XDG |
|  | LY-CoV1404<sup>36</sup> | 7MMO |
|  | C135<sup>33</sup> | 7K8Z |
|  | C110<sup>33</sup> | 7K8V |
|  | CoV2-2130<sup>14</sup> | 7L7E |
| 4 | S2H97<sup>34</sup> | 7M7W |
|  | S2X259<sup>37</sup> | 7M7W |
|  | S2X35<sup>34</sup> | 7R6W |

### 0.2 *In-Silico* Predictions

#### System preparation

For simulations, we strived to match the S-RBD region used in DMS experiments (N331-T531). For the S2H13 system, only a shorter RBD segment (S443-N501) was available. For systems C144 and C121, the Ab interacted with more than one S-RBD monomer in the crystal structure. To avoid potential loss of important interaction contributions, we included the complete spike trimer for these systems. For system C121, the structure contained two Abs interacting with S-RBD in two different conformations, hence we duplicated that system for both states and included the spike trimer: “up” state (1 RBD “up” and 1 RBD “down” in contact with the Ab) and “down” state (2 RBDs “down” in contact with the Ab). Considering one duplicated system, our analysis uncovered a total of 20 systems. System C002 required the reorientation of the CYS480 sidechain to properly form the disulfide bond with CYS488 in the RBD. Missing residues were added with the OpenMM-based PDBFixer^38^ application.

#### MD simulations

Each Ab/Ag complex was placed separately in a water box with physiological ion concentration and then energy minimization, equilibration, and 150 ns-long MD production were performed using GROMACS 2022.5 software^39,40^. Topologies were created using the parameters from the CHARMM36 force field^41^. Each system was placed in a dodecahedron simulation box such that all Ab/Ag atoms were at least 1.2 nm from the box edges and the box was then filled with TIP3P^42^ water molecules. Each box was neutralized by adding the appropriate number of Cl^−^ and Na^+^ counter-ions at a physiological concentration of 0.15 M. Systems were subsequently energy minimized under periodic boundary conditions using the steepest decent algorithm^43^. An initial equilibration phase of 1 ns was conducted under NVT conditions to facilitate the equilibration of water molecules around the proteins. The equilibration phase began at 100 K, with a linear temperature increase, concluding at 310 K. Subsequently, a second equilibration phase was performed by switching on pressure coupling, and then simulated under NPT conditions for 1 ns at a reference pressure of 1 bar, using Parrinello-Rahman coupling. Position restraints with a force constant of 1000 kJ mol^−1^ nm^−2^ were imposed during equilibration phases on all heavy atoms. Finally, production simulations were carried out under the NPT ensemble at 310 K and 1 bar each lasting 150 ns with a 2 fs timestep. For all simulation stages, periodic boundary conditions were applied and the LINCS^44^ algorithm constrained hydrogen bonds to their ideal lengths. Temperature control utilized the V-rescale^45^ option. Electrostatic interactions were computed using Particle Mesh Ewald with a real-space cutoff of 1 nm. Van der Waals interactions were computed using a cutoff of 1 nm. For systems where the S-RBD was taken into account, 100 conformers were extracted from the last 100 ns simulations every 1 ns. In systems requiring the inclusion of a complete S-trimer, we selected 10 representative structures from MD simulations through conformational clustering using the Gromos algorithm in order to reduce the computational load.^46^.

#### FoldX calculation

Snapshots from MD simulations were used as input structures for binding affinity (Δ*G*_bind_) calculations by first optimizing their geometry with the FoldX 5.0^47^ force field and six repeated energy minimizations using the *RepairPDB* command. The binding affinity between the wild-type (wt) Ab and Ag (Δ*G*_wt_) was determined using the *AnalyseComplex* command. Next, we introduced point mutations to generate different variants using the *BuildModel* subroutine. We then computed the binding affinity of the mutated complex (Δ*G*_mut_) with *AnalyseComplex* and calculated ΔΔ*G*_bind_ = Δ*G*_mut_ − Δ*G*_wt_. In the results section, we report the averages of ΔΔ*G*_bind_ or Δ*G*_bind_, calculated from either 100 individual snapshots or, in the case of trimers, 10 snapshots. The scanning region of interacting surface residues for point mutations was the same among Abs from the same class whenever possible. It includes a union set of residues on the S-RBD within 10 Å of any Abs chains in each crystal structure. It’s important to note that yeast-displayed DMS experiments encompassed a comprehensive scan of all RBD sites, including sites that are distant from the interacting region, and mutants with significantly reduced ACE2 binding or expression rates were removed from the experiments. By contrast, our approach focused on sites located within a 10 Å proximity to any Ab chain, and we conducted site saturation mutagenesis only on these selected sites. For instance, in the case of system AZD8895, 1950 escape fraction data points were reported, 1615 binding affinities were calculated, and the intersection of these datasets yielded 888 mutations for comparison analysis.

### 0.3 Comparing experimental data and predictions

The escape fraction (*f*) is not a direct measure of the binding affinity (Δ*G*_bind_), however, a mutation with a large Δ*G*_bind_ value (i.e., low affinity) has a higher chance to escape^26^. Assuming that the experiment is performed under equilibrium conditions, the binding affinity is related to the experimental escape fraction through the Hill relation^48,49^ as follows:

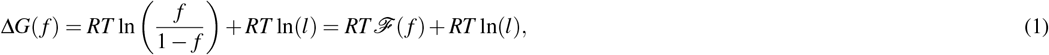

Where: *f* is the experimental escape fraction, *R* is the gas constant, *T* is the temperature in Kelvin, and *l* is the free Ab. As shown *ℱ*( *f*) is proportional to Δ*G* and is defined as a function of the escape fraction as follows:

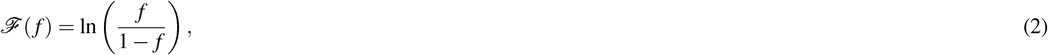

In our study, we compared predicted binding affinities and experimental escape fractions by determining the correlation between predicted Δ*G*_bind_ and *ℱ*( *f*) (Equation (2)). To assess the degree of correlation we employed two widely used statistical measures: the Pearson and Spearman correlation coefficients. These coefficients provide insights into the linear (Pearson) and monotonic (Spearman) relationships between the predicted and experimental values. Hence, a strong Pearson correlation would suggest our Δ*G*_bind_ calculations are accurately predicting *ℱ*( *f*). A strong Spearman correlation would suggest that the rank order of calculated values aligns with the rank order of experimental values.

### 0.4 Optimization of the affinity cutoff

To enhance predictive accuracy, we fine-tuned the “affinity cutoff”, that distinguishes between mutations likely to cause escape due to binding disruption from those less likely to do so. In general, a binary classifier requires a cutoff to separate positive from negative outcomes. For actual outcomes versus predicted outcomes two cutoffs are employed, a vertical and a horizontal, respectively (refer to Figure 2c). The vertical cutoff classifies the experimental data, and the horizontal cutoff classifies the predictions. These cutoffs segment the diagram into four quadrants in a confusion matrix: the top right quadrant is for True Positives (TP), where both predicted and actual outcomes are positive (i.e., both simulation and experiment agree that it is an escape mutation); the bottom right quadrant is for False Negatives (FN), where the actual outcome is positive, but the prediction is negative (i.e., the simulation predicts it is not an escape mutation, but it actually is); the top left quadrant is for False Positives (FP), where the prediction is positive but the actual outcome is negative (i.e., the simulation predicts escape mutation, but it is actually not); and the bottom left quadrant is for True Negatives (TN), where both predicted and actual outcomes are negative (i.e., both simulation and experiment agree that it is not an escape mutation).

**Figure 2.**
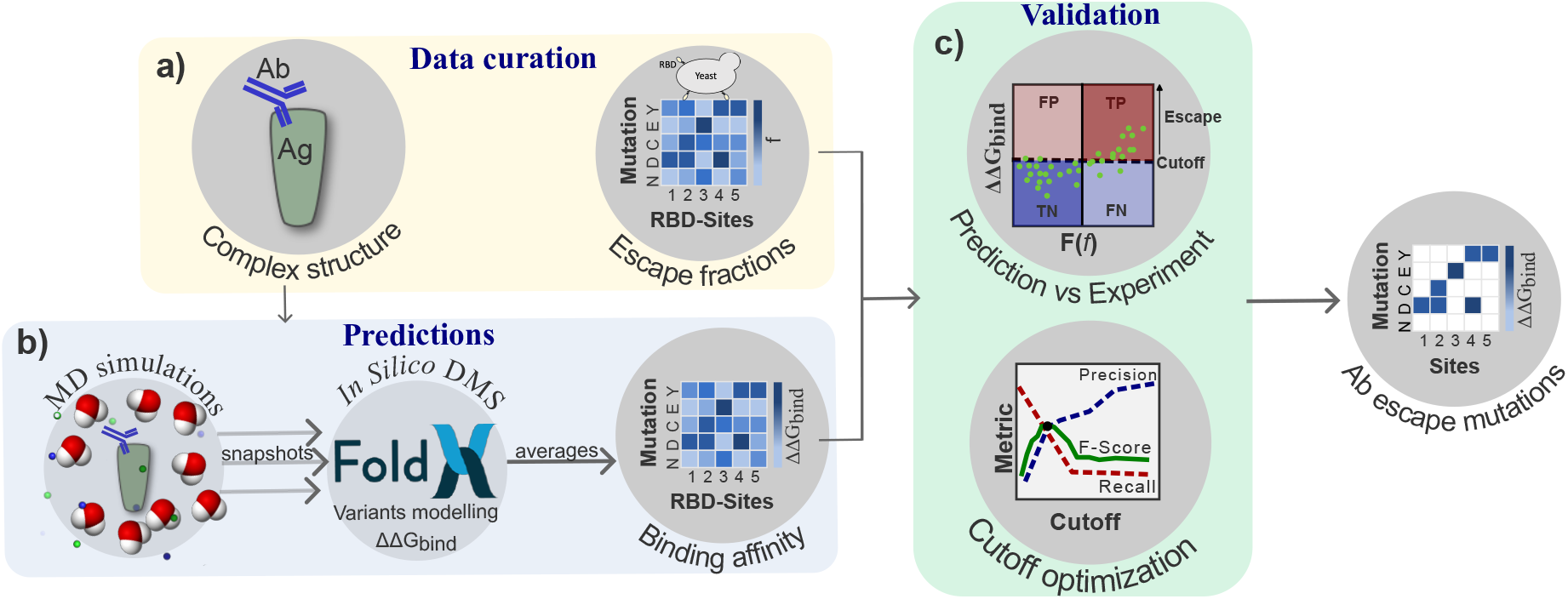
The methodology pipeline of the study is structured as follows: a) systems are selected based on the availability of data; b) binding affinity predictions are generated for each system using the MD+FoldX approach; and c) predictions are compared with experimental escape fraction data, leading to the optimization of the energy cutoff and performance evaluation, followed by the creation of an Ab-escape mutation list using the optimized cutoff.

In this study, the most important metric is TP since correctly predicting escape mutations is the goal of the method. We consider FP important as well, since it is a measure of inaccuracies in our method, but secondary to TP. Both TN and FN are less important due to the fact that Ab-escape can happen for many reasons other than Δ*G*_bind_ changes. Said another way, the goal is to predict escape, not the absence of escape. To assess the ability of MD+FoldX to correctly identify escape mutations while minimizing false predictions, we calculated the proportion of correctly identified escape mutations among all positive predictions:

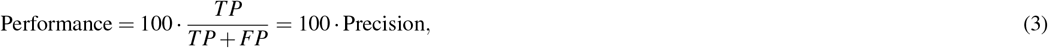

We also investigated the importance of the binding affinity cutoff in the performance of MD+FoldX. We fixed the vertical cutoff at 0.5 consistent with previous studies^11,14^ (that is, 50% of cells were found in the Ab-escape bin), and on fine-tuned the horizontal cutoff that classifies our MD+FoldX predictions as escape mutations or not. We will refer to this horizontal cutoff as the “affinity cutoff” or simply the “cutoff”. This cutoff differentiates between mutations that are more likely to lead to escape due to binding disruption (above the cutoff) and those that are less likely to cause escape (below the cutoff). To optimize the cutoff, we aimed to maximize the precision, that is, the fraction of positive outcomes correctly predicted (Equation(4)), while also maintaining a reasonable number of positive predictions. An algorithm that solely focuses on optimizing precision runs the risk of selecting a few points with very high precision but potentially missing broader relevant data. Here, we optimized the cutoff by maximizing the F-Score, a weighted average of the precision and recall^50^.

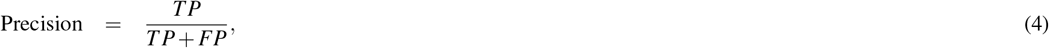

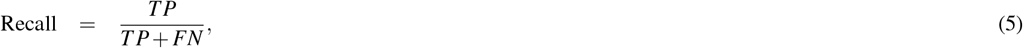

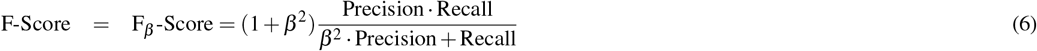

where *β* represents the weight of recall compared to precision. For our study, *β* = 0.5 was chosen meaning that precision is considered twice as important as recall (note that typical values of *β* are 0.5, 1.0, or 2.0). Precision and F-Score metrics are typically recommended for analyzing performance in unbalanced datasets, such as in our study^50^. For each system, we implemented an algorithm that iteratively tested a range of cutoffs, from the highest Δ*G* value down to Δ*G*_wt_, incrementing by 0.1 kcal/mol. The cutoff that yielded the highest F-Score was selected as the optimal value for each system. Note that we are not considering cutoffs below Δ*G*_wt_ since we seek mutations that disrupt binding compared to wild-type. For the remainder of the manuscript, we will refer to the “classic cutoff” as a value of 2.0 kcal/mol from our prior studies^21,29,51,52^, and the “optimized cutoff” as determined by maximizing the F-Score.

## Results

The purpose of this study was to evaluate the ability of the MD+FoldX method to predict antibody escape mutations using large-scale data from DMS experiments. We selected a diverse set of 19 Ab/Ag complexes targeting the SARS-CoV-2 RBD, each with structural and escape fraction data from DMS experiments. Binding affinities (Δ*G*_bind_) were predicted for each system using MD+FoldX, and Pearson and Spearman correlations to experimental escape (*ℱ*( *f*), Equation (2)) were determined. We then optimized a system-specific binding affinity cutoff based on the experimental DMS data. Subsequently, we evaluated the accuracy of the method by assessing whether the predicted escape mutations were correctly identified according to the experimental DMS data. Finally, we illustrate how the MD+FoldX method identifies escape mutations in a given system and present the list of potential escape mutations generated by our method.

The 19 Ab/Ag complexes used in our study are categorized into four distinct classes^11^ (Figure 1b and Table 1), plus an additional duplicated system for C121. Our dataset features antibodies ranging in size from 230 to 459 residues with an antigen of approximately 250 residues. When utilized, trimers consisted of over 3,300 residues. The simulation dataset contained 33763 single amino acid mutations, 4,443 of which were reported. The reported mutations are the ones reported in the empirical data set, hence they are considered to be functional in the experiments^13^. The distribution of experimental escape fraction values for all Ab/Ag complexes is depicted in Figure S1 of the Supplemental Material. The majority of experimental escape fraction values cluster around 0.0, with a secondary smaller cluster near 1.0. On average, the escape fraction value for all systems is 0.07, with at least 80% of the values falling below 0.12. The distribution of predicted ΔΔ*G*_bind_ values for all Ab/Ag complexes is as follows (Figure S2 of Supplemental Material): on average, the ΔΔ*G*_bind_ value for all systems is 0.23 kcal/mol, with 30% of the data points less than 0 kcal/mol, 60% between 0 - 1.4 kcal/mol and 10% of the data above 1.4 kcal/mol.

Figure 3 shows the Pearson and Spearman correlation coefficients between predicted Δ*G*_bind_ values and *ℱ*( *f*) (Equation (2)). Overall, we observe positive Pearson and Spearman correlation between both variables, implying that weaker binding (more positive Δ*G*_bind_) is associated with higher experimental *ℱ*( *f*). Specifically, the Pearson correlation predominantly displayed moderate to strong correlations across most systems, with a mean Pearson correlation of 0.51. Among the systems, 25% exhibited strong correlations (Pearson > 0.6), 55% displayed moderate correlations (0.6 > Pearson > 0.4), and the remaining 20% indicated weak correlations (0.4 > Pearson > 0.2). Conversely, the Spearman correlation generally showed weaker associations, with a mean Spearman correlation coefficient of 0.24. Of the systems, 10% exhibited moderate correlations (0.6 > Spearman > 0.4), while 55% displayed weak correlations (0.4 > Spearman > 0.2), and the remaining 35% indicated very weak correlations (Spearman < 0.2). Notably, in our study, the LY-CoV555 system demonstrated the strongest correlation coefficient across all systems, with Pearson’s *r* = 0.68 and Spearman’s *s* = 0.41. Conversely, the S2H13 system displayed the lowest Spearman correlation, while the C105 system exhibited the lowest Pearson correlation coefficient. Notably, the S2H14 and C121up systems showed closer correlation coefficients between Pearson and Spearman.

**Figure 3.**
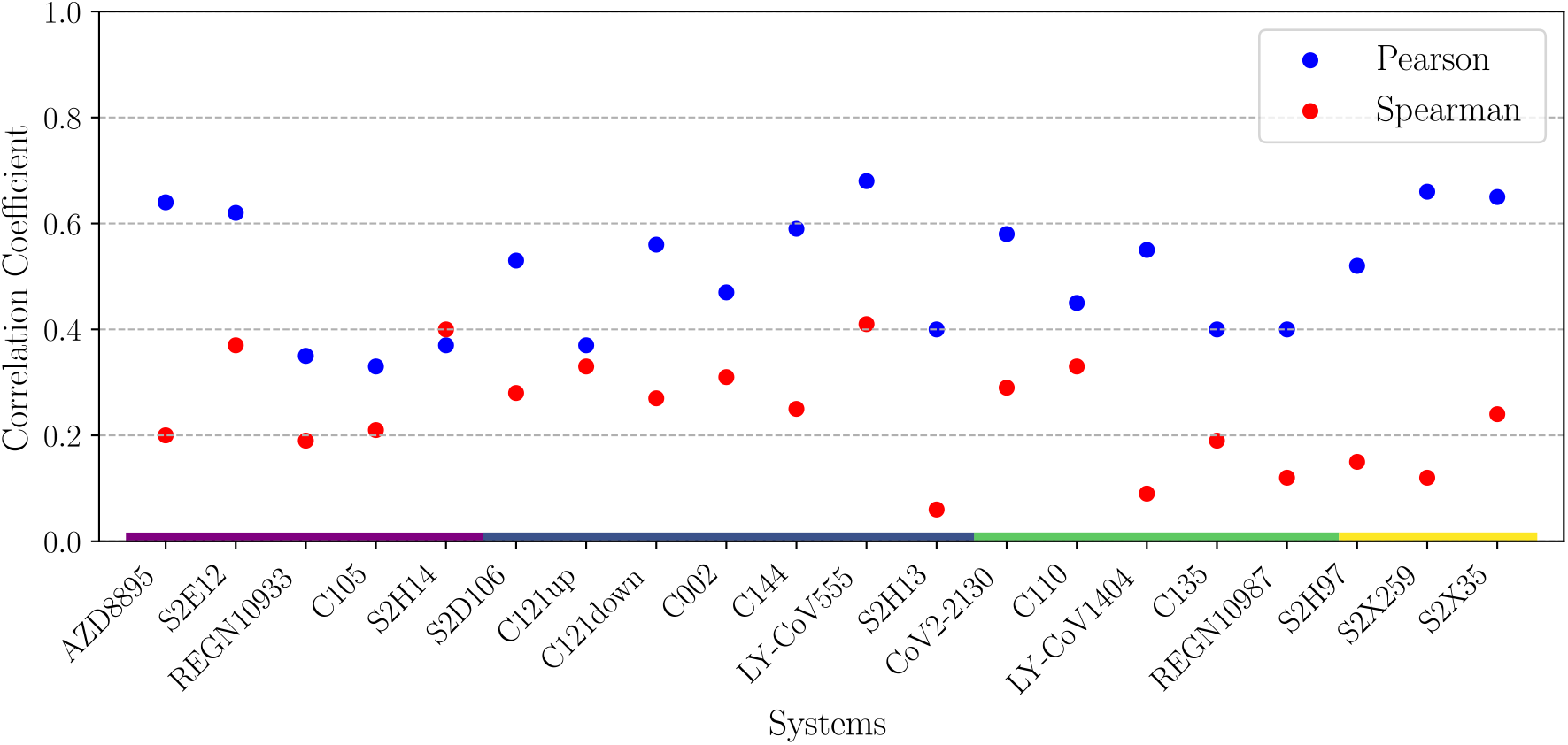
Pearson and Spearman correlation coefficients between predicted Δ*G*_bind_ values and experimental escape fractions (*f*), represented by *ℱ*( *f*) as defined in Equation (2). We show all the Ab/Ag complexes spanning the 4 Ab classes: Class 1 (purple), Class 2 (blue), Class 3 (green), and Class 4 (yellow).

In the MD+FoldX approach, the selection of affinity-driven escape mutations relies on the chosen affinity cutoff. In our previous studies, a lack of large-scale data impeded the verification of this cutoff, compelling us to adopt a cutoff of ΔΔ*G*_bind_ = 2.0 kcal/mol, here referred to as the “classic cutoff”. This approach yielded promising results in our prior studies of CB6 and B38 Abs, enabling the identification of mutations previously observed in clinically emerging variants^11^. Now equipped with extensive data, we are able to test our classic cutoff across a broad variety of systems. As part of this test, we optimized the cutoff to balance the precision and recall by maximizing the F-Score metric (6)); a reasonable approach for handling imbalanced datasets^50^. Figure 4 illustrates our optimization process using the S2H14 system as an example. Figure 4a portrays the precision, recall, and F-Score values obtained across a range of cutoffs. The optimal cutoff of -14.0 kcal/mol was chosen to maximize the F-Score. At this cutoff value, the corresponding precision is 0.55, recall is 0.59, and the F-Score reaches 0.56. Figure 4b shows the predicted Δ*G*_bind_ values against experimental *ℱ* (*f*) values. A comparison between the classic cutoff (represented by the orange dashed line) and the optimized cutoff (indicated by the green dashed line) reveals that the optimized cutoff is smaller than the classic one in this case, encompassing more data points above the cutoff. Detailed results for class 1 systems using either the classic or optimized cutoff are summarized in Table 2. Overall, a better balance between precision and recall is observed across all cases, as demonstrated by the increased F-Score when employing an optimized cutoff, compared to the classic cutoff. Systems S2H14 and C105 experienced significant performance increases using the optimized cutoff, capturing more data points as true positives and mitigating false negatives.

**Table 2.** Example of performance metrics of Class 1 systems by utilizing the classic and optimized horizontal cutoff. Cutoff_Δ*G*_ and Cutoff_ΔΔ*G*_ correspond to cutoffs expressed in terms of Δ*G* or ΔΔ*G*, respectively. TP is true positive, FP is false positive, TN is true negative, and FN is false negative.

| Classic Cutoff |  |  |  |  |  |  |  |  |  |  |
| --- | --- | --- | --- | --- | --- | --- | --- | --- | --- | --- |
| | System | TP | FP | TN | FN | Precision | Recall | F-Score | $\text{Cutoff}_{\Delta G}$ | $\text{Cutoff}_{\Delta\Delta G}$ |
| Class 1 | AZD8895 | 22 | 10 | 838 | 19 | 0.69 | 0.54 | 0.65 | -12.7 | 2.0 |
|  | S2E12 | 27 | 7 | 817 | 38 | 0.79 | 0.42 | 0.67 | -11.2 | 2.0 |
|  | REGN10933 | 19 | 26 | 798 | 44 | 0.42 | 0.30 | 0.39 | -17.5 | 2.0 |
|  | C105 | 3 | 6 | 816 | 64 | 0.33 | 0.04 | 0.15 | -9.5 | 2.0 |
|  | S2H14 | 12 | 11 | 781 | 85 | 0.52 | 0.12 | 0.32 | -12.8 | 2.0 |
| Optimized Cutoff |  |  |  |  |  |  |  |  |  |  |
| | System | TP | FP | TN | FN | Precision | Recall | F-Score | $\text{Cutoff}_{\Delta G}$ | $\text{Cutoff}_{\Delta\Delta G}$ |
| Class 1 | AZD8895 | 20 | 6 | 842 | 21 | 0.77 | 0.49 | 0.69 | -12.5 | 2.2 |
|  | S2E12 | 34 | 7 | 817 | 31 | 0.83 | 0.52 | 0.74 | -11.5 | 1.7 |
|  | REGN10933 | 16 | 17 | 807 | 47 | 0.48 | 0.25 | 0.41 | -16.9 | 2.6 |
|  | C105 | 45 | 64 | 758 | 22 | 0.41 | 0.67 | 0.45 | -11.1 | 0.4 |
|  | S2H14 | 57 | 46 | 746 | 40 | 0.55 | 0.59 | 0.56 | -14.0 | 0.8 |

**Figure 4.**
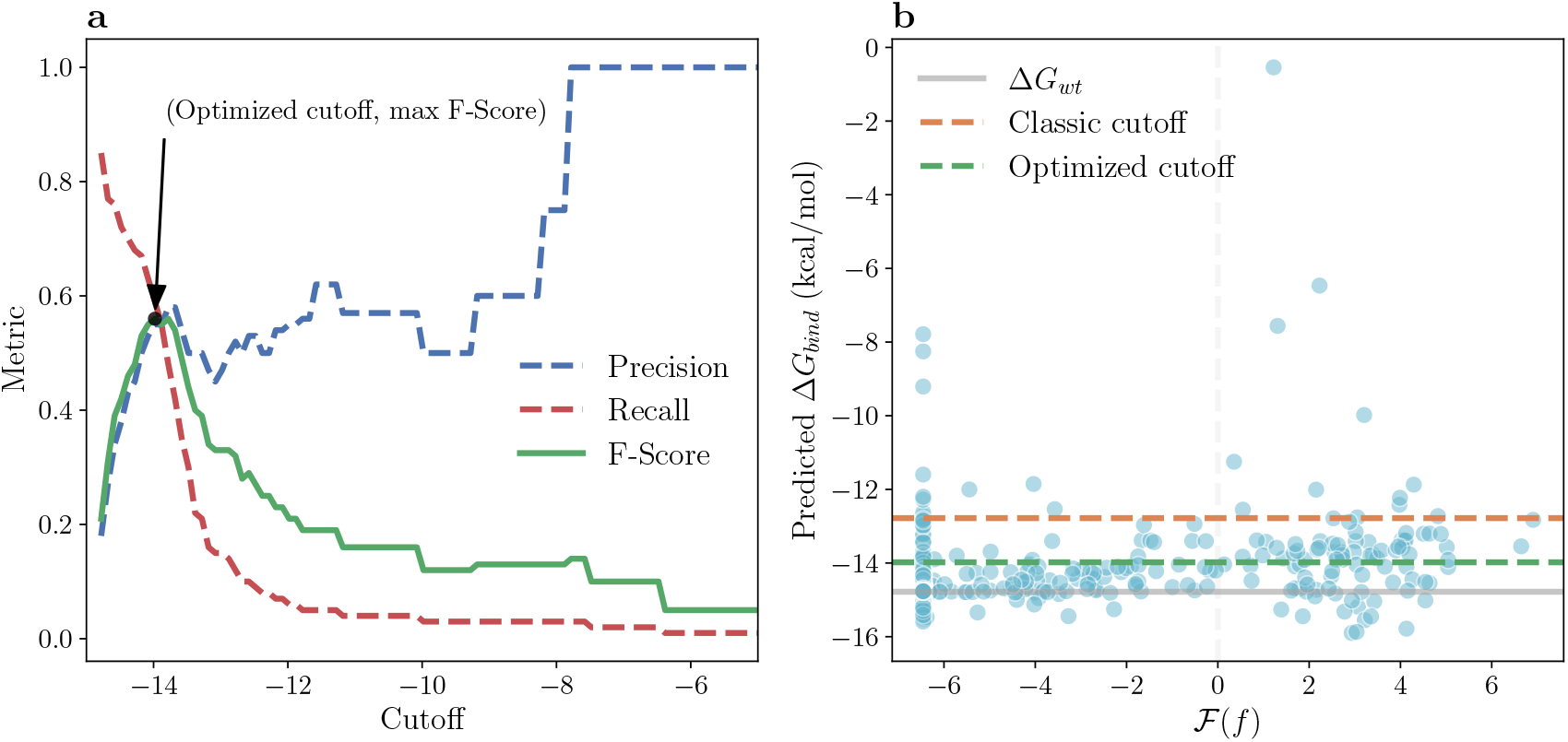
Example of a cutoff optimization using the S2H14 Class 1 system. a) Precision, recall, and F-Score values across a range of cutoffs. b) Scatter of predicted Δ*G*_bind_ values against *ℱ* (*f*); the green dashed line represents the optimized cutoff, the orange dashed line indicates the classic cutoff, the grey dashed line represents the escape fraction value at which the 50% of the cells expressing a specific variant escape Ab binding, and the solid grey line indicates the value of the Δ*G*_bind_ for the wild-type Ag.

Table 3 shows that the median optimized cutoff for all the systems is 1.8 kcal/mol, with an interquartile range (IQR) of 1.8, indicating variability in the cutoff values across the Ab classes. Table 3 illustrate that each Ab class presents a distinct range of cutoff values. Class 4 has the highest median cutoff at 2.9 kcal/mol, while Class 3 has the lowest at 0.8 kcal/mol. The IQR of 1.0 for Class 4 and 0.8 for Class 3 indicates a relatively narrow spread of values. Figure 5 illustrates the cutoff values derived from the optimization process for the complete set of systems. Among the systems, 14 exhibit cutoffs within the range of 1.0 kcal/mol to 3.0 kcal/mol, 5 systems have values below 1.0 kcal/mol, and 1 system is above 3.0 kcal/mol.

**Table 3.** Quartiles and Interquartile Range for all systems and per Class.

| Class | Q1 | Median | Q3 | IQR |
| --- | --- | --- | --- | --- |
| Class 1 | 0.8 | 1.7 | 2.2 | 1.4 |
| Class 2 | 1.5 | 2.6 | 2.8 | 1.3 |
| Class 3 | 0.4 | 0.8 | 1.2 | 0.8 |
| Class 4 | 2.5 | 2.9 | 3.5 | 1.0 |
| All | 0.8 | 1.8 | 2.6 | 1.8 |

**Figure 5.**
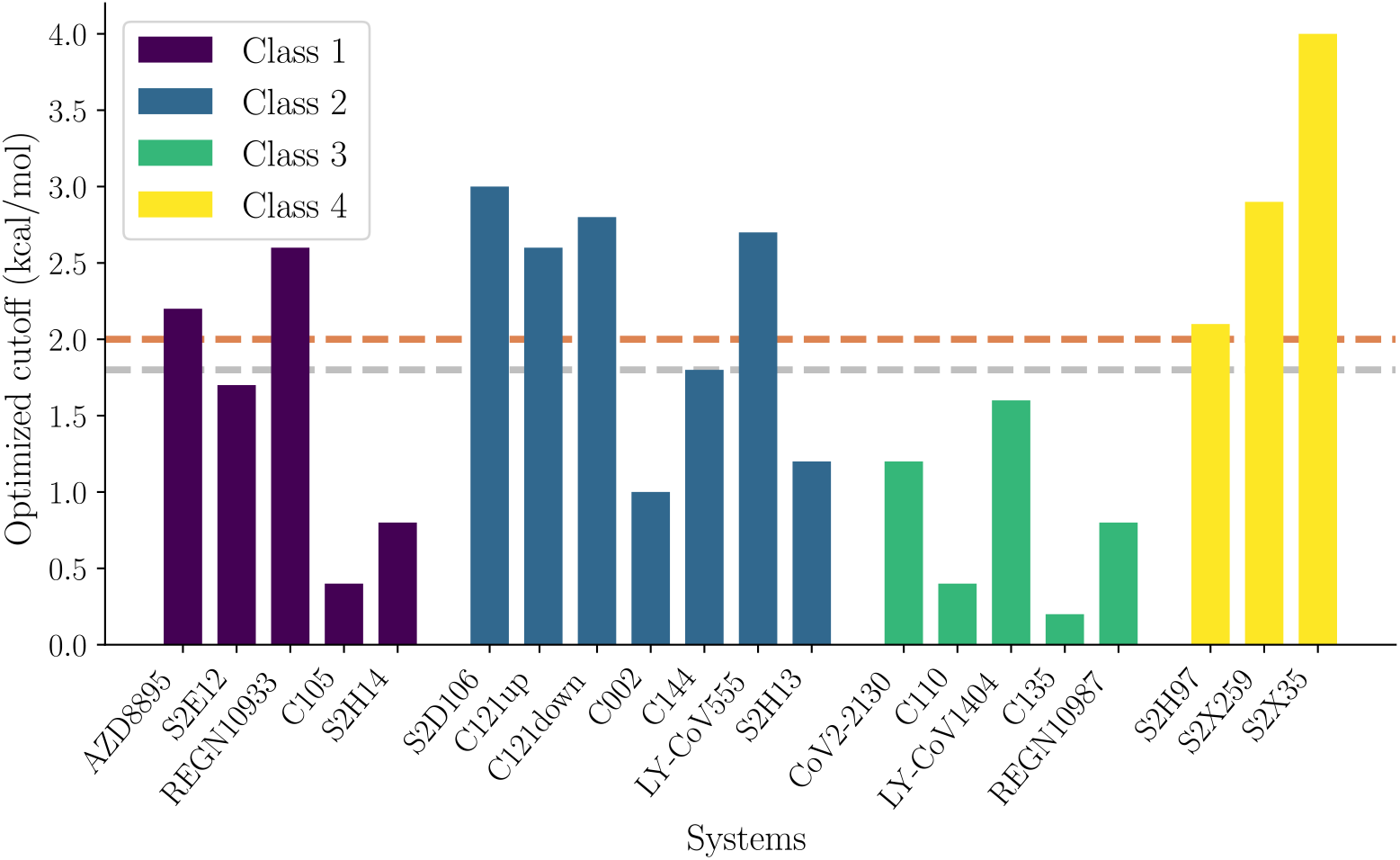
Optimized cutoff_ΔΔ*G*_ for each system. The orange dashed line represents the classic cutoff and the grey dashed line the median optimized cutoff_ΔΔ*G*_.

Figure 6 illustrates the performance achieved when utilizing the optimized cutoffs. The mean overall systems was 66%. It can be observed that 70% of the systems exhibit a performance above 50%. Notably, the systems S2X35 and S2D106 demonstrate the highest performance, while C135 and S2H97 display the lowest performance. In contrast, the mean performance achieved when utilizing the classic cutoff was 59%.

**Figure 6.**
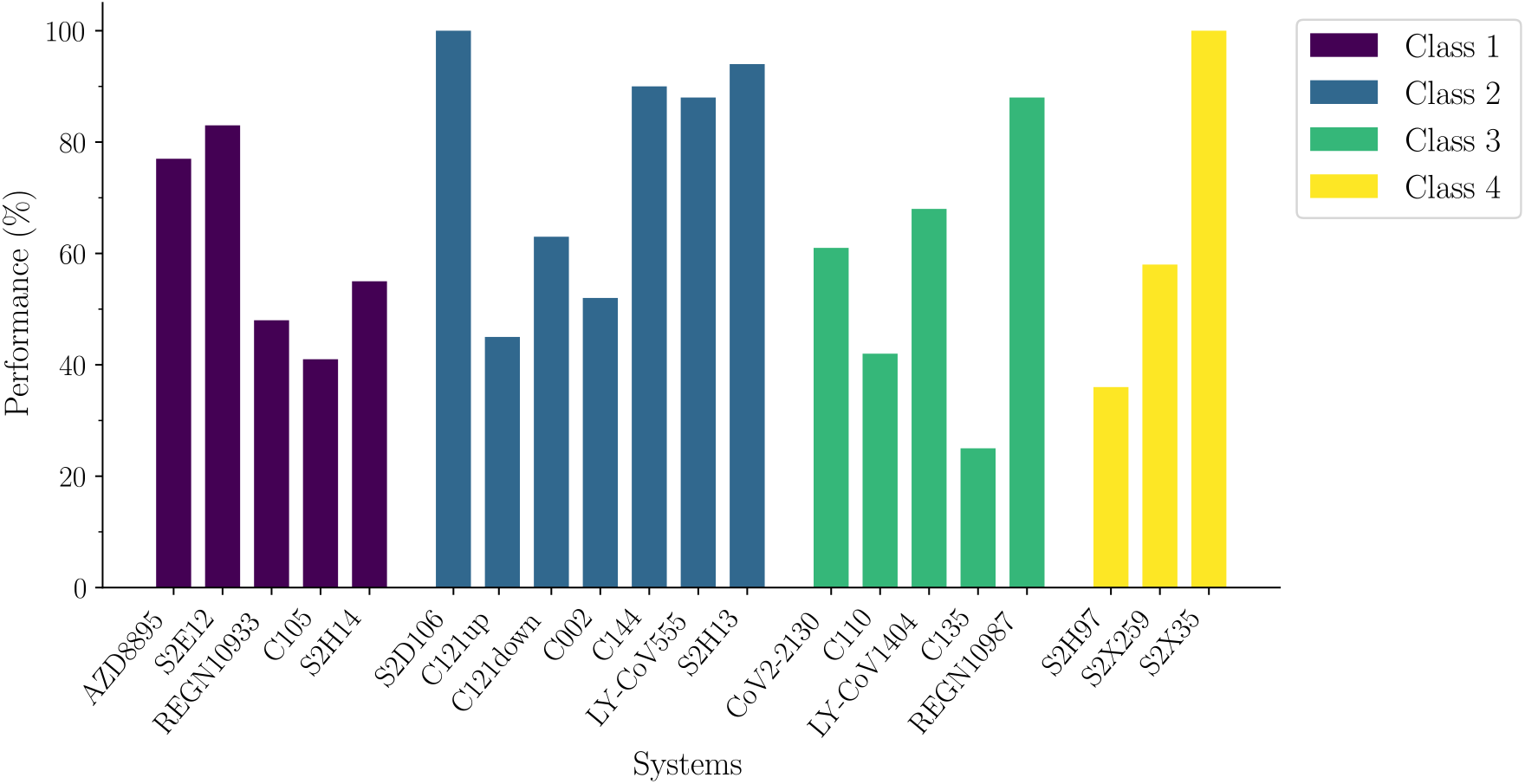
Performance comparison across systems as defined by Equation (3) using optimized cutoffs for each system.

As an example of how escape mutations are identified by the MD+FoldX approach with the optimized cutoff, consider system LY-CoV555. This Ab classified as Class 2 Ab, has been granted an emergency use authorization for the treatment of COVID-19^35^. Figure 7 shows the predicted ΔΔ*G*_*bind*_ for each point mutation within this system. The darker squares on the heatmap indicate a higher positive ΔΔ*G*_*bind*_, signifying mutations that significantly weaken the binding affinity between the Ag and Ab, hence increasing the likelihood of escape. A detailed enumeration of escape mutations about all the systems under study is available in Tables S1-S4 of the Supplemental Material.

**Figure 7.**
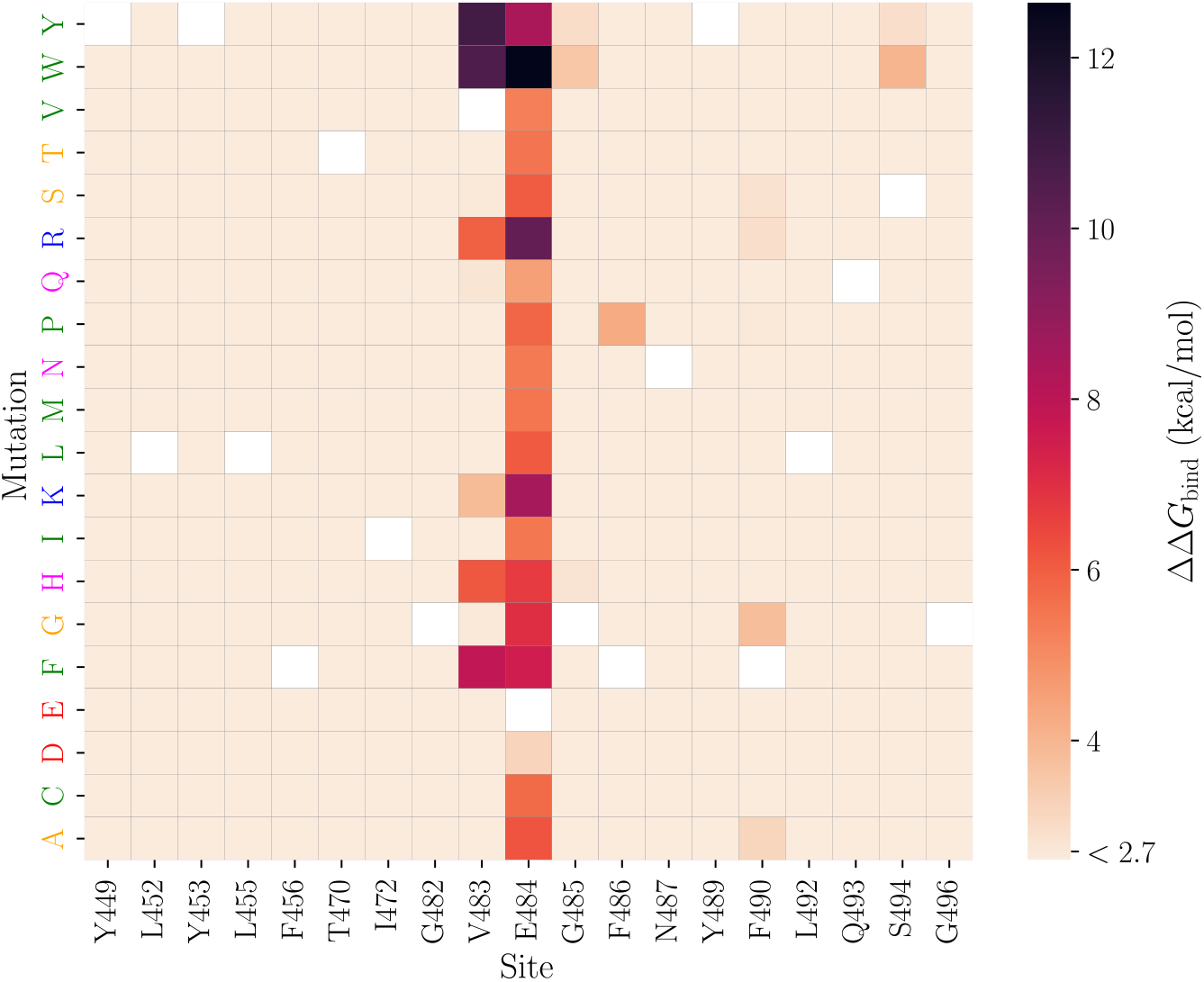
Heat map of predicted ΔΔ*G*_*bind*_ values using MD+FoldX for mutations in the RBD when in complex with LY-CoV555 Class 2 Ab. The horizontal axis displays the amino acid positions in the RBD along with the corresponding native amino acids. The vertical axis represents the mutation of amino acids colored according to physicochemical properties: small non-polar, hydrophobic, polar, negatively charged and positively charged. White squares represent the amino acids of the wild-type. For clarity in visualization, sites where all mutations led to changes in binding free energy of less than 0.5 kcal/mol were omitted from the heat map.

## Discussion

In this study, we evaluated the ability of the MD+FoldX method to predict SARS-CoV-2 RBD escape mutations using a comprehensive deep mutational scanning dataset. We focused on 19 Ab/Ag systems with structural information of the complexes and escape fractions from the yeast-based DMS technique. Our analysis revealed a positive correlation between predicted affinity and experimental escape fraction. We found that fine-tuning predicted affinity cutoffs using empirical data allowed for more accurate escape mutation identification, underscoring the importance of tailoring cutoffs to specific Ab/Ag interactions rather than adopting a one-size-fits-all approach. Our results demonstrate the potential of MD+FoldX to identify significant affinity-driven escape mutations, some of them already present in variants of concern and interest, and also highlight the need for the development of more accurate methods.

This study includes 19 Ab/Ag systems across four distinct Ab classes, all directed against the SARS-CoV-2 S-RBD. This dataset covers Abs targeting similar epitopes, i.e. Abs within the same class, as well as those targeting different structural epitopes, i.e. Abs from different classes. Analysis of the escape fraction data for all the systems showed that only a small proportion of mutations exhibit escape (high escape fractions), consistent with the prevailing understanding that the majority of mutations have negligible effects on antigenicity^53^. In addition, our analysis of binding affinity predictions for all systems showed that the majority of the values were destabilizing (ΔΔ*G* > 0 kcal/mol); this is consistent with previous studies in a series of globular proteins using the FoldX algorithm where approximately 70% of mutations were destabilizing^54^ and with experiments where random mutations tend to be destabilizing^55^. Notably, our investigation encompasses a substantial dataset comprising 4,443 single amino acid mutations across the 19 antibody-antigen systems, each accompanied by experimental escape fraction data. In comparison, related studies have examined significantly fewer mutations: Tandiana and coworkers validated 27 mutations in different antibody/hen egg white Lysozyme complexes with experimental data^56^, Gonzalez and coworkers validated 253 mutations in Ab/Ag complexes^27^, Miller and coworkers validated 114 mutations in 27 Ab/Ag systems^29^, Beach and coworkers validated 8 mutation in Ab/Ag systems^30^, and Sharma and coworkers validated 22 mutations in Ab/Ag systems^57^. Validating *in silico* predictions of affinity-driven antibody escape mutations remains challenging due to the scarcity of large-scale experimental data, like because obtaining binding affinity data for a large number of mutations is costly and time consuming. However, our research shows that the extensive escape fraction data from Bloom’s large-scale studies^34–36^ provides a promising alternative for validating this type of prediction.

One primary obstacle we faced was devising a suitable approach to accurately link binding affinity with escape fractions. Here, we assume that the experiment was performed under equilibrium conditions and hence the binding affinity is related to the experimental escape fraction through the Hill relation^48,49^. We found that the Pearson and Spearman correlation coefficients were consistently positive via the Hill relation. Our mean Pearson correlation coefficient of 0.51 is comparable to the accuracy observed in previous MD+FoldX studies, where a correlation coefficient of 0.46 was reported for predicted versus experimental ΔΔ*G*_bind_^27^. In contrast, our mean Spearman correlation coefficient was 0.24, also consistent with the limited performance of other similar methods studying Ab/Ag systems, where Spearman correlation coefficients rarely exceed 0.28^27,58^.

While Δ*G*_bind_ serves as a valuable metric for assessing binding strength, its correlation with the escape fraction may be influenced by a multitude of complex factors inherent to the binding process: 1) Δ*G*_bind_ provides a thermodynamic view of affinity, but escape fractions also include kinetic factors such as binding and unbinding rates. Thus, while high Δ*G*_bind_ coupled with low kinetic barriers could result in high escape fractions (indicating a strong correlation), the presence of high kinetic barriers alongside high Δ*G*_bind_ can obscure this relationship due to impeded escape despite weak binding^59,60^. 2) Protein folding variations can significantly affect epitope accessibility and stability. Altered epitopes due to folding changes can reduce antibody affinity, thereby affecting the likelihood of escape. In particular, the yeast-DMS approach excludes mutants with significant misfolding^33^. Conversely, predictive models often do not calculate antigen folding stability; when they do, their accuracy can be variable, posing additional challenges to accurately predicting escape mutations^19–23^. 3) The presence of multimeric structures, such as trimers in spike proteins, could introduce cooperative binding effects that could affect binding strength and escape. In yeast-DMS experiments, antibodies bind to monomeric yeast-expressed RBDs, and therefore cannot fully capture mutational effects on spike-trimer conformation or effects on antibodies that bind quaternary epitopes^33^. 4) Differences in the glycosylation patterns of yeasts compared to human cells may influence antibody recognition of glycan-rich epitopes. In particular, N-linked glycans on yeast-expressed proteins are more mannose-rich than those on mammalian-expressed proteins^33^. In this study, our simulations do not include glycans and hence our prediction accuracy would suffer for such an epitope. 5) Finally, variations in the structural dynamics of the Ab/Ag complexes, the influence of solvent effects like bridging water molecules, and the presence of allosteric effects or cooperative binding events, are potentially significant factors that are very challenging to account for in simulations. Indeed, all of the factors listed above underscore the challenges of directly correlating binding energies with escape fractions and highlight the intricate interplay between structural, kinetic, and biological phenomena in antibody-antigen interactions.

Given the factors listed in the previous paragraph that could impact the correlation between predictions and escape fraction data, we will consider three distinct cases. Case 1: Antibodies bind to monomeric non-glycan epitopes. Here, Δ*G*_bind_ is expected to correlate with *f* via the Hill relation (Equation (1)), meaning that significantly larger Δ*G*_bind_ in mutants compared to wildtype lead to higher likelihood of escape. Case 2: Antibodies bind glycan-free epitopes that span several monomeric units. In this case, the correlation between Δ*G*_bind_ and escape fractions *f* may not be strong since the experimental setup includes only monomeric spike protein leading to larger escape fractions (*f* > 0.5). Case 3: Antibodies bind glycan-rich epitopes. In this case, Δ*G*_bind_ may not correlate well with escape fractions *f* since yeast cells produce different glycans. In our study, systems are predominantly consistent with Case 1, including AZD8895, S2E12, S2H14, REGN10933, C105, LY-CoV555, S2D106, S2H13, REGN10987, LY-CoV1404, CoV2-2130, S2H97, S2X259, C002, and S2X35. Here, Δ*G*_bind_ could be expected to correlate with *f* via the Hill relation. Positive high Pearson correlation coefficients associated with these systems (Pearson = 0.28 − 0.68) indicate that larger Δ*G*_bind_ are associated with high escape fractions. By contrast, C121 is consistent with Case 2, Abs targeting glycan-free quaternary epitopes^11^. In this system the spike trimer structure has 2 Abs bound, the first one (C121up) interacts primarily with 1 RBD in the “down” conformation but also has minor contact with 1 RBD in the “up” conformation. A second Ab (C121down) interacts primarily with 1 RBD in the “down” conformation and has minor contact with a second RBD in the “down” conformation. Although the correlation between Δ*G*_bind_ and escape fractions *f* are not expected to be consistent, Δ*G*_bind_ values could still serve as useful indicators of potential escape events. Systems C110 and C135 are consistent with Case 3, both featuring a glycosylated epitope containing an N343 glycan^11^. Finally, the C144 system is a hybrid between cases 2 and 3; its epitope spans multiple monomers and includes a glycosylated site (N434). However, its primary interaction is with one of the RBDs that is not glycosylated, and may explain the reasonable correlation found for this system.

In this study, we found that adjusting predicted affinity cutoffs based on empirical data was crucial for accurately identifying escape mutations. Overall, there was significant variability in the optimized cutoff values across different systems and classes. This suggests that the optimal cutoff for predicting escape mutations varies depending on the specific Ab/Ag system, indicating that a one-size-fits-all approach may not be suitable for all systems and highlighting the complexity and specificity of Ab/Ag interactions. The challenge of selecting an appropriate cutoff value is further emphasized by the varied approaches used in different studies to pinpoint SARS-CoV-2 immune hot spots. Chang et al.^20^ used a ΔΔ*G* = 0 kcal/mol cutoff to classify binding stability, with negative values indicating stabilization and positive ones indicating destabilization. Huang et al.^19^ developed an immune-escaping score by merging binding free energy measurements with variant frequency, using variants of concern and interest to determine mutation frequency cutoffs. Mauria et al.^22^ refined binding affinity and escape fraction cutoffs using receiver operating characteristic curve analysis, setting ΔΔ*G* = −0.7 kcal/mol and an escape fraction of 0.001753. In this study, we fixed the escape fraction cutoff and concentrated on fine-tuning the Δ*G*_bind_ horizontal affinity cutoff. As observed in our results, significant variability among the systems was found, however, the distribution of optimized cutoffs tends to center around our classic cutoff of 2.0 kcal/mol showing that it could serve as a reasonable starting point for initial predictions. It may be particularly applicable in the absence of system-specific data to guide the optimization of the cutoff value. The data reveals Class 3 Abs possess a minimal tolerance for energy changes, corresponding to lower escape cutoffs, whereas Class 4 Abs exhibit a higher tolerance, requiring more substantial energy changes to allow mutations to escape.

It is crucial to weigh the clinical implications of the cutoff selection carefully: an overly stringent cutoff may overlook potential escape mutations, whereas too lenient a cutoff might overpredict escape, potentially triggering unnecessary concern or interventions. Hence, the optimized cutoff must ensure that predictions are both accurate and practically useful for guiding therapeutic strategies and vaccine design. Aiming for the ambitious goal of predicting escape mutations solely through Δ*G* values, adopting a proposed stepwise approach could prove instrumental. We propose the use of system-dependant optimized cutoff whenever possible. If data for optimizations of the particular system is lacking, we can formulate three distinct lists per class based on varying cutoff values to cater to different scenarios. For an optimistic approach, a cutoff based on Q1 in Table 3 is recommended. This threshold may encompass some non-escape mutations but is aimed at generating a concise list suitable for preliminary investigations. The second approach adopts a cutoff based on the median, designed to offer balanced results, making it appropriate for a wide range of cases. Lastly, a more stringent cutoff based on Q3 in Table 3 is suggested for scenarios prioritizing the identification of highly probable escape mutations. While this may not capture all potential escape mutations, it is particularly valuable in therapeutic antibody development, where the focus is on mutations with significant implications for treatment efficacy. In scenarios where systems diverge from those evaluated in this study and lack tailored data for cutoff optimization, the established classic cutoff of 2.0 kcal/mol may provide a practical baseline for initial predictions.

In our analysis, we emphasized positive results, since accurate prediction of escape cases is crucial. To evaluate the ability of the method to correctly identify positive cases (mutations that are more likely to be escape mutations due to binding disruption) while considering possible errors in the method (false positives), we compute the positive predictive value. This is the proportion of correctly identified escape mutations out of all positive predictions. For Case 1 systems, high performance is possible, provided that the Δ*G*_bind_ predictions are accurate, as this is the case where Abs binding glycan-free monomeric epitope as in the experiments. In Case 2 systems, Δ*G*_bind_ values could still serve as useful indicators of potential escape and high performance is possible, assuming accurate Δ*G*_bind_ predictions. In Case 3 systems, however, Δ*G*_bind_ is likely to be inaccurate, resulting in low escape prediction performance. Overall, precision-based performance varied across the systems: seven exhibited high performance (Performance > 80%), fourteen showed moderate performance (Performance > 50%), and six had low performance (Performance < 50%). This highlights the method’s effectiveness in predicting escape mutations but also emphasizes the need for more accurate and efficient binding-free energy estimation methods. Systems classified under Case 1 showed performance ranging from 23% to 100%. Case 2 systems had a performance range of 45% to 90%. As expected, Case 3 systems displayed relatively low performance in predicting escape mutations, ranging from 25% to 43%.

In general, our method was able to identify significant variants of concern and interest such as R346K, L452R, E484K/A/Q, F486F/V, K417N/T, and N501Y. Specifically, the escape mutations E484K/A/Q, identified using the MD+FoldX methodology for LY-CoV555 system, were documented across multiple SARS-CoV-2 variants of concern, including the Alpha (B.1.1.7), Beta (B.1.351), Eta (B.1.525), Mu (B.1.621), and Omicron BA.1 and BA.2 (B.1.1.529) lineages, as well as the Kappa (B.1.627.1) lineages^16^. Similarly, the mutation F486P was observed in the Omicron (XBB.1.5) subvariant^61^. E484K that substitutes a positively charged residue with a negatively charged one was demonstrated to escape LY-CoV555 Ab reducing neutralization^35^. Interestingly, S2X259, a broad sarbecovirus neutralizer, exhibits an escape profile confined to a single mutation, G504D, as revealed by DMS and *in vitro* escape selection experiments^37^. The fact that we were able to predict these mutations suggests that our approach has potential to uncover key resistance mechanisms that could undermine current antibody-based treatments and vaccines. This insight provides valuable data for ongoing SARS-CoV-2 research, particularly in developing effective therapeutic antibodies and improving future vaccine formulations. Moreover, understanding these mutation patterns helps researchers monitor viral evolution, ultimately contributing to global efforts in preventing and managing COVID-19 and similar infectious diseases.

Future studies should explore how interactions between mutations, known as epistatic effects, could cause non-additive impacts on binding affinity and immune escape. This research would be crucial for understanding the emergence of antibody escape mutations in clinical variants that contain multiple mutations. In addition, our study suggests that there is a need for methods to more accurately estimate binding free energy changes upon mutation at a reasonable computational speed. Furthermore, the impact of antibody structural variability and flexibility on computational predictions warrants further investigation, especially given the challenges posed by lower-resolution structures. Although FoldX provides more accurate free energies for high-resolution crystal structures (less than 2.6 Å)^31^, its effectiveness with cryo-EM-derived and low-resolution crystal structures requires thorough evaluation. While affinity plays a key role in antibody escape, it does not encompass all aspects of escape potential. Machine learning methods could significantly enhance the detection of escape mutations. These models trained using escape fractions from DMS experiments, could incorporate additional relevant features beyond mere binding affinity.

## 1 Conclusions

In this study, we demonstrated the potential of the MD+FoldX method for predicting antibody escape mutations by leveraging deep mutational scanning data from various antibody-antigen complexes targeting the SARS-CoV-2 RBD. The insights gained here were only possible due to the extensive data generated by DMS in several studies^14,32–37^, emphasizing the crucial role of ongoing data collection and analysis in advancing infectious disease research. Our study showed a positive correlation between predicted binding affinity changes and experimental escape fractions, suggesting that MD+FoldX can predict escape mutations in some cases. The strength of the correlation was system-dependent. It is important to acknowledge that even with perfect calculations, Δ*G* will not perfectly correlate to escape fractions due to other factors such as binding rates, protein folding, multimeric epitopes, and glycosylation differences. Yet, it is also the case that a sufficiently large change in Δ*G* will invariably result in escape. Our results emphasize that the ideal cutoff for predicting escape mutations should be tailored to the specific Ab/Ag interaction, challenging the efficacy of a universal standard and highlighting the complexity of Ab/Ag interactions. We recommend a stepwise decision-making approach using the cutoffs found in this work as a reference for similar systems. Overall, our study shows that MD+FoldX can streamline the prioritization of mutations for in-depth analysis, and ultimately facilitate detection of potential escape mutations. However, it also underscores that there is still a need for more accurate methods for estimating binding free energy changes upon mutation at a reasonable computational speed.

## Acknowledgements

This project is supported by the P3-R1 Grant Matching Program at the University of Idaho (UI). NIH grant P20GM104420 supported this research; views expressed are solely those of the authors. Research reported in this publication was supported by an Institutional Development Award (IDeA) from the National Institute of General Medical Sciences of the National Institutes of Health under grant number P30GM103324. We acknowledge the use of the Falcon supercomputer resources^62^. The funders had no role in study design, data collection and analysis, decision to publish, or preparation of the manuscript.

## Author contributions statement

LAC, JSP and FMY conceived and conceptualized the project. FMY and JSP supervised the project. LAC and JEB conducted the simulations. The results were analyzed by FMY, JSP, LAC, and JEB. The initial draft of the manuscript was prepared by LAC and JEB, and all authors contributed to reviewing and editing the manuscript.

## Availability of data and materials

Code, inputs, and raw data to reproduce plots for this work are available on GitHub at https://github.com/YtrebergLab/SARS-CoV-2_Ab-escape.

