## Supplemental Material for "Exploring the ability of the MD+FoldX method to predict SARS-CoV-2 antibody escape mutations using large-scale data"

### 1 Assessing data distribution

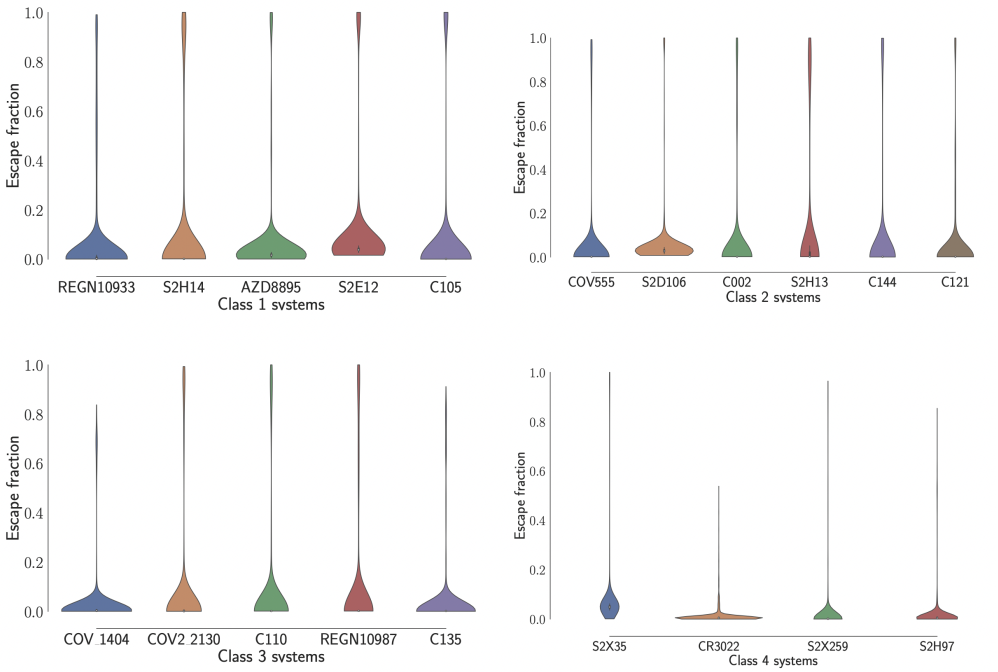


Figure S1: Escape fraction ($f$) distribution per system and class.

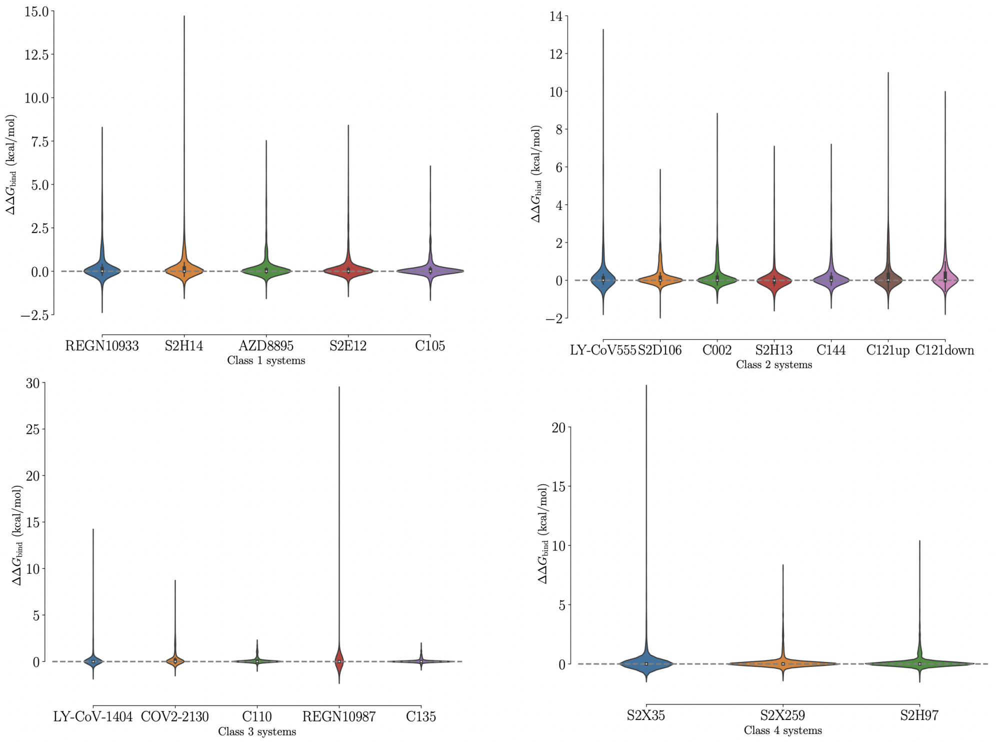


Figure S2: $\Delta\Delta G_{Bind}$ distribution per system and class.

#### 2 Escape lists

Table S1: Class 1 escape mutation list per system and site generated using the optimized cutoffs. Shown in bold font are sites and mutations that have been previously observed in VOC, VOI or their sub-variants.

| System | Site | Mutation |
|---|---|---|
| AZD8895 | **F486** | **S**, H, K, G, T, E, **P**, D, A, N, Q, R, W, C |
| | N487 | Y, F |
| S2E12 | A475 | N, K, D, E |
| | G476 | R, K, H, N, Q, E, D, T |
| | **S477** | P |
| | **T478** | D |
| | **F486** | **P**, D, A, N, Q, R, **V**, W, C, **S**, Y, H, K, T, G, E |
| | N487 | H, F, Y, E |
| REGN10933 | **F486** | G, A, E, N, P, Q, K, C, D, T, R, S, V |
| | N487 | F, Y |
| | **Q493** | Y |
| C105 | **K417** | M, L, I, C, **N**, A, V, G, **T**, Q, P, F, E, S, D |
| | D420 | R, K |
| | Y453 | V, E, T, Q, C, N, V, I, S, A |
| | L455 | P, C, A, E, T, G, H, N, S |
| | F456 | C, I, G, V, T, Q, A |
| | A475 | R, Y |
| S2H14 | R403 | A, S |
| | K417 | E |
| | Y449 | F, Q, W, S, D, E, M, T, N, L, K, C, H, V, I, Q |
| | L455 | S, W, A, E, G, H |
| | F456 | T, A, G, K |
| | **G496** | I, E, V, D |
| | **Q498** | G, D, E, W |
| | T500 | E, D, Y, W |
| | **N501** | M, H, F, P, C, I, Q, V, G, **Y**, L, A, T, S, W, R |

Table S2: Class 2 escape mutation list per system and site generated using the optimized cutoffs. Shown in bold font are sites and mutations that have been previously observed in VOC, VOI or their sub-variants.

| System | Site | Mutation |
|---|---|---|
| S2D106 | **E484** | W, Y |
| | **F486** | **P** |
| C121up | **E484** | **A**, F, **K**, S, Y, R, W, H, G |
| C121down | **E484** | G, N, V, F, H, D, P, T, **A**, W, C, I, K, R, S, Y, M |
| C002 | **L452** | K, **R**, W, V, F |
| | I472 | E, G, D, W |
| | Y473 | A |
| | V483 | K, S, D |
| | **E484** | I, V, C, G, W, T, S, **A** |
| | G485 | T, R, P |
| | **F486** | R, L, G, Q, D, E, **P**, K, T, W, N, I, **V** |
| | **Q493** | Y, A, D, W, C, T, F, I, V |
| | S494 | R |
| C144 | **E484** | L, Y, N, P, I, H, V, C, **Q**, T, W, G, F, **K**, S, R, M, **A** |
| | **F486** | G |
| LY-CoV555 | V483 | Q, Y, R, K |
| | **E484** | **K**, R, S, M, **A**, L, Y, N, P, I, V, H, D, C, **Q**, F, W, T |
| | G485 | H |
| | **F486** | **P**, G, A, R, **S** |
| | S494 | Y, W |
| S2H13 | V483 | T, E, N, Y, K, R, A, S, H, D, Q, F, G, W |
| | G485 | S, A, N, E, M, P, C, H, Q, K, R, T, D |
| | **F486** | **S**, G, D, E, **P**, T |

Table S3: Class 3 escape mutation list per system and site generated using the optimized cutoffs. Shown in bold font are sites and mutations that have been previously observed in VOC, VOI or their sub-variants.

| System | Site | Mutation |
|---|---|---|
| CoV2-2130 | **R346** | W, F, C, S, D, L, A, G, E, Q, T, V, I, N |
| | K444 | N, E, L, H, W, D, I, A, R, G, V, P, F, Y, C, S, T |
| | V445 | P, G |
| | G446 | L, P, I, M, P, T, E, D, H, Q, Y, N, L, F, V |
| | N448 | Y |
| | Y449 | D, K, R |
| | S494 | K, R |
| C110 | **R346** | D, E |
| | K444 | D, A, G |
| | G447 | D, E |
| | Y449 | K, S, A, Q, R, T, D, P, V, E |
| | **L452** | H, E, T, K, S, W, D, Y |
| | F490 | D, E, Q, K, A, N, R, P, S |
| LY-CoV-1404 | K444 | E, D, I, A, G, V, P, F, S, T |
| | V445 | R, K, Y, W, D, N, H, G, F, S |
| | G446 | P |
| | P499 | W, F, H, R, K, Y |
| C135 | **R346** | Y, D, **K**, A, G, H, E, Q, T, V, I, N, F, W, C, S |
| | N440 | K, L, Y |
| | L441 | S |
| | K444 | D, I, P, E |
| REGN10987 | V445 | S, G, A, H |
| | G446 | I, M, F, W, Q, R, C, L, V, A, D, S, H, T, Y, E, K, P, N |

Table S4: Class 4 escape mutation list per system and site generated using the optimized cutoffs.

| System | Site | Mutation |
|---|---|---|
| S2H97 | K462 | D, E |
| | S514 | Y, R |
| | E516 | T, R |
| | L518 | W, A, S |
| S2X259 | P384 | K |
| | R408 | E |
| | G504 | D, H, T, Y, P, E, L, I, V |
| S2X35 | T376 | F, Y, H |
| | G404 | K, Y, W, M, E, Q, I, K, L, F, R, D, H |
